# Antiviral RNAi response against the insect-specific Agua Salud alphavirus

**DOI:** 10.1101/2021.12.20.473494

**Authors:** Mine Altinli, Mayke Leggewie, Marlis Badusche, Rashwita Gyanwali, Christina Scherer, Jonny Schulze, Vattipally B. Sreenu, Marvin Fegebank, Bernhard Zibrat, Janina Fuss, Sandra Junglen, Esther Schnettler

## Abstract

Arboviruses transmitted by mosquitoes are responsible for the death of millions of people each year. In addition to arboviruses, many insect-specific viruses (ISVs) have been discovered in mosquitoes in the last decade. ISVs, in contrast to arboviruses transmitted by mosquitoes to vertebrates, cannot replicate in vertebrate cells even when they are evolutionarily closely related to arboviruses. The alphavirus genus includes many arboviruses, although only a few ISVs have been discovered from this genus so far. Here, we investigate the interactions of a recently isolated insect-specific alphavirus, Agua-Salud alphavirus (ASALV), with its mosquito host.

RNAi is one of the essential antiviral responses against arboviruses, although there is little knowledge on the interactions of RNAi with ISVs. Through knock-down of transcripts of the different key RNAi pathway (siRNA, miRNA and piRNA) proteins, we show the antiviral role of *Ago2* (siRNA), *Ago1* (miRNA), and *Piwi4* proteins against ASALV in *Aedes aegypti* derived cells. ASALV replication increased in *Dicer2* and *Ago2* knock-out cells, confirming the antiviral role of the siRNA pathway. In infected cells, mainly ASALV-specific siRNAs are produced while piRNAs, with the characteristic nucleotide bias resulting from ping-pong amplification, are only produced in *Dicer2* knock-out cells. Taken together, ASALV interactions with the mosquito RNAi response differs from arthropod-borne alphaviruses in some aspects, although they also share some commonalities. Further research is needed to understand whether the identified differences can be generalised to other insect-specific alphaviruses.

## Introduction

Mosquitoes are efficient vectors for many medically important arthropod-borne viruses (arboviruses) from several RNA virus families such as *Flaviviridae, Togaviridae, Bunyavirales, Reoviridae*, and *Rhabdoviridae* (Weaver & Reisen, 2010). Arboviruses have a complex life cycle consisting of replication in both vertebrate and invertebrate hosts. In the last decade, many viruses that are restricted to invertebrate hosts (*i*.*e*. that cannot replicate in vertebrate hosts) have also been discovered (Atoni et al., 2019). These viruses, generally termed insect-specific viruses (ISVs), have been discovered from all major arbovirus families. They are considered promising for many applications, from vaccine development to arbovirus transmission control tools (Agboli et al., 2019). Nevertheless, our knowledge of many important aspects of ISV biology is limited, such as their interactions with the vector species they infect (Altinli et al., 2021).

Arboviruses establish asymptomatic persistent infections in mosquito vectors which are attributed to the efficiency of the mosquito innate immune system. As a part of the mosquito innate immune system, RNA interference (RNAi) pathways play a major role in regulating arbovirus infections (Donald et al., 2012; Leggewie & Schnettler, 2018). There are three RNAi pathways in mosquitoes: micro (mi)RNA, small interfering (si)RNA and P-element induced wimpy testis (PIWI)-interacting (pi)RNA pathways (Donald et al., 2012; Leggewie & Schnettler, 2018). The siRNA pathway is triggered by double-stranded (ds)RNA and categorised as exogenous or endogenous depending on the origin of the dsRNA. Among these, the exogenous (exo-)siRNA pathway is considered the primary antiviral defence mechanism for mosquitoes and other insects (Bronkhorst & Van Rij, 2014; Leggewie & Schnettler, 2018). The exo-siRNA pathway can be induced by dsRNA derived either from viral replication or RNA secondary structures, which are cut by *Dicer2* (*Dcr2*) into virus-derived siRNAs (vsiRNA), that are 21 nucleotides in length (Donald et al., 2012; Leggewie & Schnettler, 2018). These vsiRNAs are then incorporated into the RNA-induced silencing complex (RISC), specifically to the *Argonaute2* (*Ago2*) protein, and guide the complex to target complementary viral RNA for subsequent cleavage; resulting in the inhibition of virus replication. vsiRNAs specific to arboviruses are produced during infection by all major arboviruses, proving an interaction with the exo-siRNA pathway (Liu et al., 2019). Furthermore, knock-down or knock-out of key players involved in the exo-siRNA pathway, *Dcr2* and *Ago2* proteins, led to an increase in replication of all tested arboviruses, supporting the antiviral role of these proteins and the exo-siRNA pathway against arboviruses in mosquitoes (Liu et al., 2019; Scherer et al., 2021).

The miRNA pathway is known to regulate gene expressions of endogenous transcripts in various organisms, including mosquitoes. The miRNA pathway starts by cleaving primary miRNAs into precursor (pre-)miRNA molecules in the nucleus. After exportation into the cytoplasm, pre-miRNA is cut to miRNA/miRNA* duplexes of 21-22nt in size by *Dicer1*. miRNAs then guide miRISC (RISC complex associated with the miRNA pathway), including the *Ago1* protein, to degrade and/ or inhibit translation of (partially) complementary single-stranded (ss)RNAs (Asgari, 2014, 2015). However, our knowledge of the antiviral role of the miRNA pathway in mosquito-virus interactions is limited.

Arbovirus specific piRNAs of 25-29 nts length have also been reported in infected mosquitoes and mosquito-derived cells (Miesen et al., 2016; Varjak, Leggewie, et al., 2018). In *Aedes aegypti-*derived cells, virus-derived piRNA (vpiRNA) biogenesis is *Piwi5/6* (depending on the investigated virus) and *Ago3* dependent for Sindbis, Chikungunya and Dengue viruses (Miesen et al., 2015, 2016; Varjak, Dietrich, et al., 2018). The transcripts are bound by *Ago3* (sense) and *Piwi5/6* (antisense) and processed in the ping-pong amplification cycle. Resultant vpiRNAs have either a bias for uridine at position one or adenine at position 10 in the antisense and sense sequences, respectively (U1, A10) and a complementary region of 10 nucleotides (Miesen et al., 2015, 2016). In contrast, another Piwi protein, *Piwi4*, does not directly bind vpiRNAs of viral or transposon origin (Miesen et al., 2015) but preferentially binds antisense piRNAs derived from endogenous viral elements (EVEs). These EVEs can be integrated into the mosquito genome during RNA virus infection and act as an “adaptive immune response” combined with the produced vpiRNAs and *Piwi4* (Tassetto et al., 2019). The knock-down of *Piwi4* transcripts resulted in increased virus titer supporting its antiviral role. In contrast, the knock-down of piRNA pathway proteins did not have a strong antiviral role against the tested arboviruses so far (Schnettler et al., 2013; Varjak, Donald, et al., 2017; Varjak, Leggewie, et al., 2018; Varjak, Maringer, et al., 2017a); except for Rift Valley Fever virus (Dietrich, Jansen, et al., 2017). *Piwi4* has been shown to interact with proteins of the piRNA and siRNA pathways; however, its antiviral activity against an arthropod-borne alphavirus is independent of *Dcr2* activity in the Aag2 cells (Varjak, Maringer, et al., 2017a).

Compared to arboviruses, ISV interactions with the mosquito RNAi pathways are less studied. Studies on ISVs mainly focused on detection of virus specific small RNAs in persistently infected cell lines. Here, the production of vsiRNAs and, in some cases, vpiRNAs were detected for different families, including *Flaviviridae, Birnaviridae* and *Phenuiviridae* (Agboli et al., 2019; Frangeul et al., 2020; Öhlund et al., 2021) Our knowledge on RNAi-ISV interactions is further limited for insect-specific alphaviruses, as no persistently infected cell lines is known and (Blitvich & Firth, 2015; Bolling et al., 2015) only four insect-specific alphaviruses have been identified in mosquitoes so far: Eilat virus (EILV), Taï Forest alphavirus (TALV), Mwinilunga alphavirus (MWAV) and Agua-Salud Alphavirus (ASALV) (Hermanns et al., 2017, 2020; Nasar et al., 2012; Torii et al., 2018). So far, only the latter has been studied for its interactions with the RNAi response (Hermanns et al., 2020). Indeed, ASALV infection in *Aedes albopictus*-derived cells induces the production of vsiRNAs, but lack the production of vpiRNAs. Moreover, it is unknown whether the siRNA pathway acts antiviral against ASALV.

In addition to the mosquitoes’ ability to control virus replication through the RNAi pathways, viruses can also suppress the RNAi response. Indeed, some ISVs such as Culex Y virus and Mosinovirus, are known to interfere with the RNAi response (Fareh et al., 2018; Schuster et al., 2014; van Cleef et al., 2014) by encoding a RNAi suppressor protein. However, it is not known whether this is the case for insect-specific alphaviruses.

Here, we investigated the interactions of ASALV with the mosquito RNAi pathways in detail. We show the antiviral role of the exo-siRNA pathway against ASALV by using *Ae. aegypti* derived *Dcr2* and *Ago2* knock-out cell lines. ASALV-specific siRNAs were still produced in the absence of *Ago2* but decreased in the *Dcr2* knock-out cell line. ASALV triggered vpiRNA production through the ping-pong production pathway only in *Dcr2* knock-out cells. By knocking down additional key RNAi transcripts, we further show the involvement of *Ago1, Ago2* and *Piwi4* as antiviral against ASALV in *Ae. aegypti* derived cells.

## Methods

### Cell lines

Aag2-AF5 (ECACC 19022601) is a single-cell clone of *Aedes aegypti* derived Aag2 cells. Aag2-AF319 (ECACC 19022602) is a *Dcr2* knock-out (KO) cell line derived from AF5 cells (Varjak, Maringer, et al., 2017b), and AF525 is an *Ago2* knock-out cell line also derived from AF5 cells (Scherer et al., 2021). *Aedes albopictus* derived C6/36 cells were used for virus production.

All cell lines were kept in Leibovitz’s L15 Medium (ThermoFisher Scientific) supplemented with 10 % tryptose phosphate broth (Gibco Life Technologies), 10 % fetal bovine serum (ThermoFisher Scientific), and 1 % penicillin-streptomycin (ThermoFisher Scientific). All cell lines were grown at 28 °C.

### ASALV stock

Previously isolated and plaque purified ASALV was used for all experiments (Hermanns et al., 2020). Virus stocks were produced by inoculating C6/36 cells. The supernatant was harvested upon observation of morphological changes and was cleared from cell debris by centrifugation. For TCID50 virus quantification, 4×10^4^ C6/36 cells/ per well were seeded in 96-well plates 2 hours before infection. Serial dilutions were performed in L15 complete media.

### dsRNA synthesis

Primers specific to *Ae. aegypti Ago1, Ago2, Ago3, Piwi4, Piwi5, Piwi6* (Schnettler et al., 2013) and LacZ (Aedes-T7-BGal F/R) (Carissimo et al., 2015) flanked by T7 RNA polymerase promoter sequences were used to amplify gene-specific fragments. Amplified fragments were validated by Sanger sequencing. PCR products were used for *in vitro* transcription and subsequent column-based purification using the MEGAscript RNAi kit (Thermo Fisher Scientific) according to the manufacturer’s instructions.

### Growth kinetics

4×10^5^ AF5 cells per well were seeded in 12-well plates a day prior to infection and kept at 28 °C overnight. Cells were infected with ASALV at a multiplicity of infection (MOI) of 0.1. After 1 hour of incubation, the infectious medium was replaced with 1 ml of fresh L15 with supplements. Samples were taken at different time points (0, 24, 48, 72 hours post-infection (hpi)). Infection and negative controls were performed in triplicates, and three independent experiments were performed. The amount of viral RNA in the supernatant was quantified using RNA isolated from supernatant samples with TRIzol LS (Invitrogen) according to manufacturer’s protocol. QuantiTect SYBR Green qRT-PCR one-step kit (Qiagen) was used to quantify ASALV with previously established primers (Hermanns et al., 2020). Samples were run in technical triplicates. An in-run calibrator and an external standard curve were used to perform an absolute quantification using Roche Light Cycler 480 II.

### ASALV infection in knock-out cells and small RNA sequencing

3×10^5^ cells/well (AF5, AF525, AF319) were seeded in 24-well plates and infected with ASALV at MOI 0.5 the following day. Total RNA of infected cells was isolated at 48 hpi with Trizol according to manufacturer’s protocol using glycogen as a carrier. QuantiTect SYBR Green qRT-PCR one-step kit (Qiagen) was used to quantify ASALV with previously established primers (Hermanns et al., 2020). ASALV RNA fold change was calculated using the 2^-ΔΔCT^ method with Ribosomal protein S7 RNA as the housekeeping gene and AF5 cells as the control group.

To investigate the production of ASALV specific small RNAs in AF5, AF525 and AF319 cells, 8×10^5^ cells were seeded in a 6-well plate and infected with ASALV (MOI 1). Total RNA was isolated at 48 hpi with TRIzol (Ambion), according to manufacturer’s protocol with glycogen as a carrier. Small RNAs of 1 µg total RNA were sequenced using BGISEQ-500 at BGI Tech (Hong Kong, China) as previously described (Scherer et al. 2021). For one of the AF525 samples (Figure S1), total RNA was sequenced at IKMB (Kiel, Germany), using 100 ng total RNA for library preparation with the NEXTFLEX® Small RNA-Seq Kit v3 (PerkinElmer Inc., Waltham, MA, USA), followed by library sequencing on one lane NovaSeq6000 SP v1.0 (2×50bp). Data analyses were performed as previously described (Varjak, Dietrich, et al., 2018). The ASALV genome sequence was used as template (MK959115). Small RNA sequencing data is available in the NCBI Sequence Read Archive under BioProject ID PRJNA725665.

### Knock-down experiments

2.5 × 10^5^ AF5 cells/well were seeded in 24-well plates the day before transfection with 200 ng of gene-specific dsRNAs or control dsRNA (dsLacZ) per well, and transfected using 1 ul of Dharmafect2 reagent (GE Dharmacon). For siRNA knock-downs in knock-out cells, 20 nM of either *Piwi4* specific siRNAs or control siRNA (Horizon Discovery) was transfected using 2 µl Dharmafect2 reagent (GE Dharmacon), as previously described (Varjak, Maringer, et al., 2017a). The following day, ASALV infection (MOI 0.5) was performed. At 48 hpi, total RNA was isolated from cells using TRIzol (Ambion). cDNA of 1.5 µg RNA was produced using M-MLV reverse transcriptase (Promega) and Oligo(dT)15 primers (ThermoFisher Scientific) according to the manufacturer’s protocol. SYBR Green quantitative RT-PCR for mRNA targets was performed using gene-specific primers (Table S1) and Ribosomal protein S7 RNA as the housekeeping gene transcript. Results were analysed using the 2^-ΔΔCT^ method with LacZ dsRNA samples as control. All qPCR reactions were performed in technical triplicates.

### RNA silencing suppressor assay

To assess whether the presence of ASALV in cells could suppress the RNA silencing response, AF5 cells were seeded in 24-well plates (1.8 × 10^5^ cell/well) one day prior to ASALV infection (MOI 10). The day following the infection, cells (ASALV or mock infected) were transfected with Firefly and Renilla luciferase expression constructs, pIZ-Fluc and pAcIE1-Rluc (Ongus, Roode, Pleij, Vlak, & van Oers, 2006; Varjak, Maringer, et al., 2017), and either 0.5 ng dsRNA (either dsFluc or dsLacZ as a negative control) or 0.1 ng siRNA (either siFluc or siHyg as a negative control) using 1 µl of Dharmafect2. 24 hours post transfection, the cells were lysed and luciferase was measured with the Dual luciferase assay (Dual Luciferase Reporter Assay system, Promega) according to manufactures protocol on a Glomax luminometer.

### Statistical analyses

R (version 3.5.2) was used for statistical analyses. First, normality (Shapiro Wilk test) and variance (F-test) of the data were tested. The student’s t-test was used for normally distributed homoscedastic data, or the Welch t-test was used for normally distributed heteroscedastic data. p<0.05 was considered as statistically significant.

## Results

### ASALV efficiently replicates in AF5 cells

The successful replication of ASALV has been previously shown in *Ae. albopictus* derived C6/36 and U4.4 cells (Hermanns et al., 2020). To verify that ASALV could replicate in *Ae. aegypti* derived AF5 cells; a cumulative growth curve was performed by collecting supernatant every 24 hours until 72 hours post-infection (hpi). The growth curves showed that ASALV efficiently replicates in AF5 cells (Figure 1) without any visible cytopathic effect.

**Figure 1:**
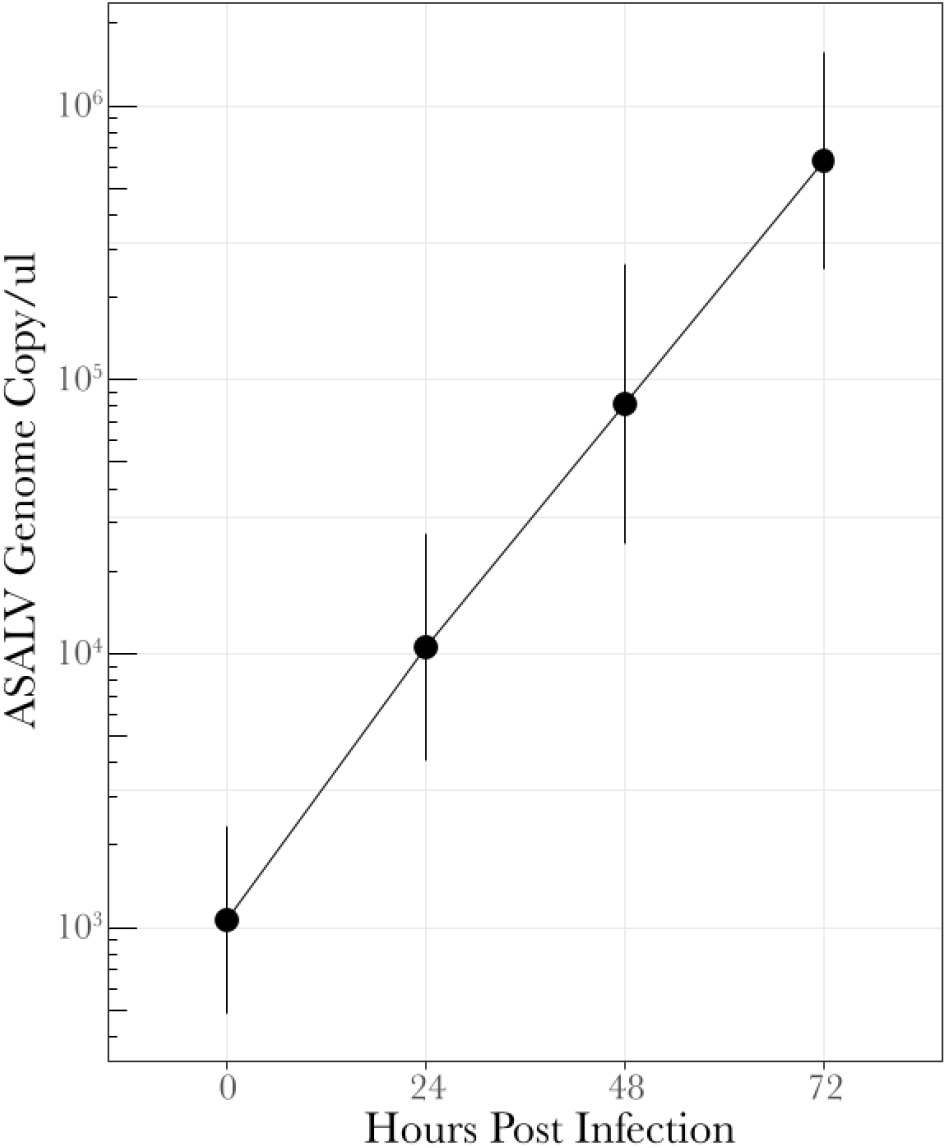
Growth Kinetics of ASALV in *Aedes aegypti* derived AF5 cells. AF5 cells were infected with ASALV with an MOI of 0.1. The supernatant was collected at different time points (0, 24, 48 and 72 hpi), and ASALV RNA was quantified by qRT-PCR. The average of three independent replicates (performed in triplicates) is shown with SEM.

### ASALV replication increases in *Dcr2* (AF319) and *Ago2* (AF525) knock-out cells

To investigate the effect of the siRNA pathway on ASALV replication, *Dcr2* (AF319) and *Ago2* (AF525) knock-out (KO) cells and control AF5 cells were infected with ASALV (MOI 0.5). ASALV RNA fold change in the KO cells compared to AF5 cells at 48 hpi was quantified by qPCR. ASALV RNA increased significantly in AF525 (t = 5.2385, df = 7, p= 0.001) and AF319 cells (t = 21.654, df = 7, p<0.001, Figure 2) compared to AF5 control cells.

**Figure 2:**
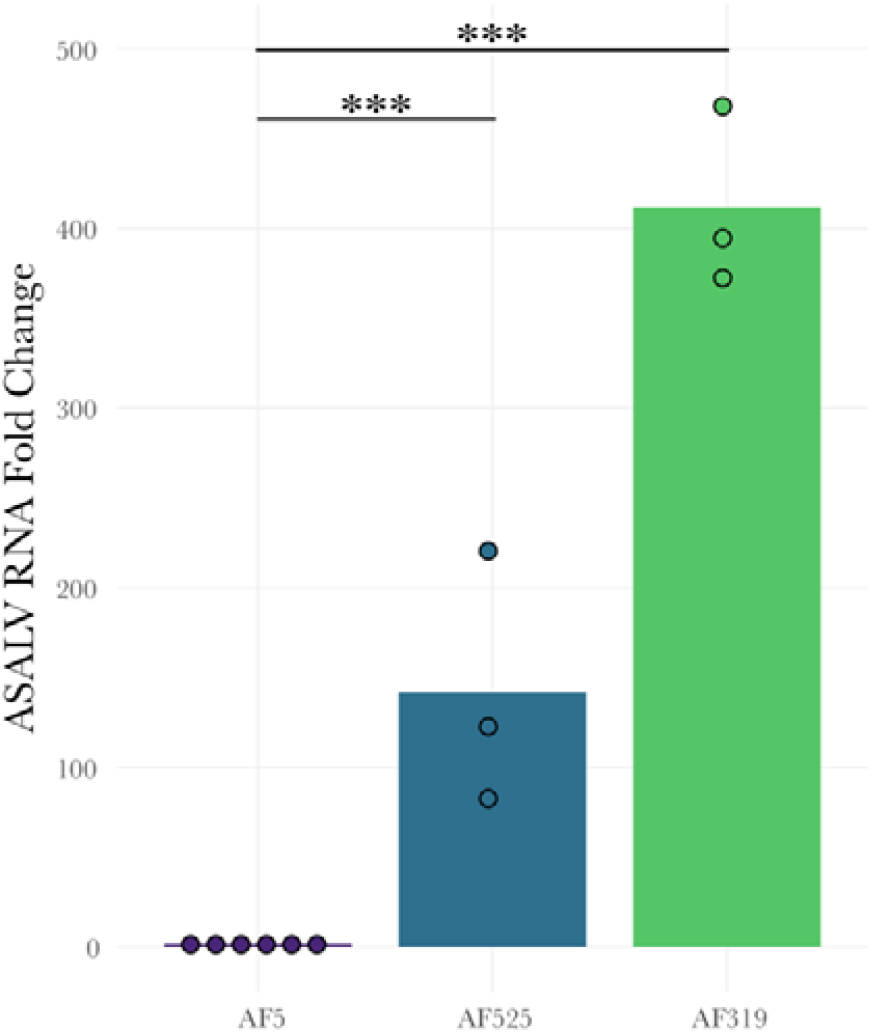
Increased ASALV replication in *Dcr2* (AF319) and *Ago2* (AF525) *Ae. aegypti* derived knock-out cells. AF319, AF525 and AF5 cells were infected with ASALV (MOI 0.5). ASALV RNA fold change in infected cells was quantified at 48 hpi, using the 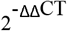 method with Ribosomal protein S7 RNA as housekeeping gene and AF5 cells as control. Three independent replicates were performed for AF525, and AF319 cells (n=3) and AF5 controls were repeated for each group (n=6). Bar plots represent the mean of the replicates that were performed (***: p<0.001).

### piRNA-sized small RNAs with ping-pong characteristics are only produced in *Dcr2* KO cells

To investigate the production of ASALV-specific small RNAs in the different cells, small RNA sequencing of ASALV infected cells was performed in *Ae. aegypti*-derived AF5, AF319 and AF525 cells. Cells were infected with ASALV (MOI 0.5), and total RNA was isolated at 48 hpi, followed by small RNA sequencing and bioinformatics analysis. Two independent replicates per cell line were performed, resulting in similar findings (Table 1 Figure 3, Figure S1-S2).

**Table 1:**
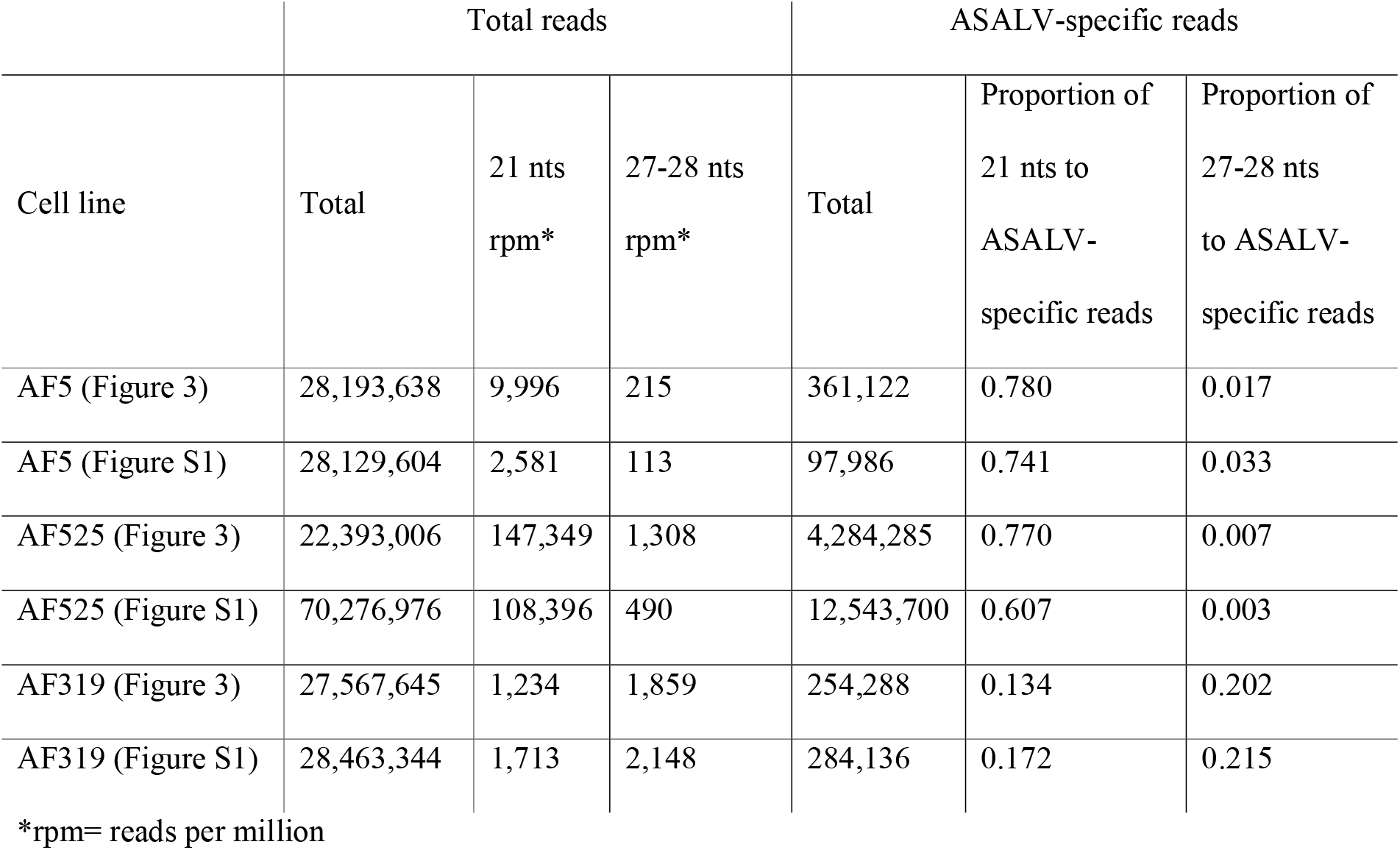
Total and ASALV-specific small RNA reads in *Ae. aegypti* derived AF5, AF525(*Ago2* KO) and AF319 (*Dcr2* KO) cells.

**Figure 3:**
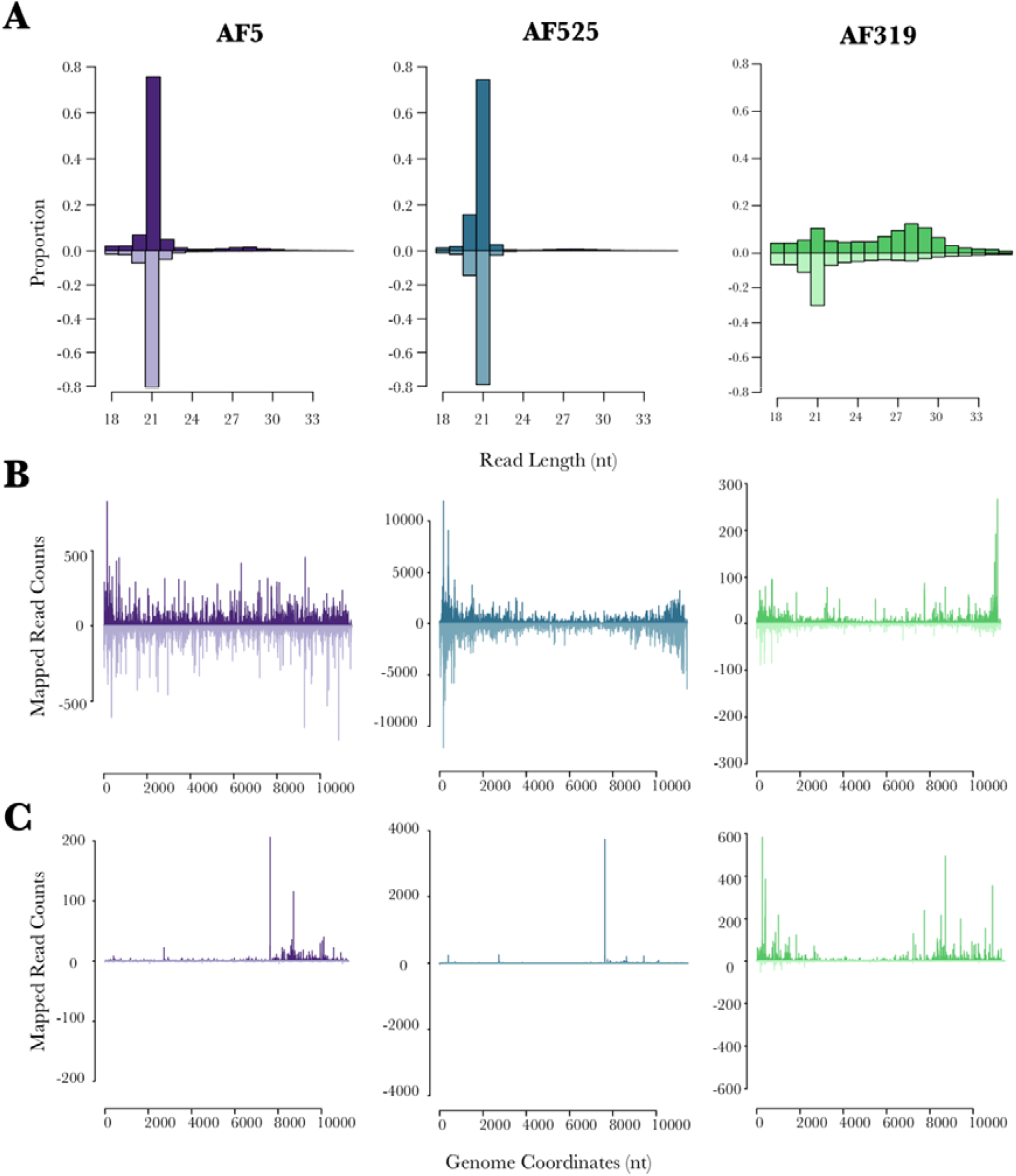
ASALV-specific small RNA production in *Ae. aegypti* derived AF5, AF319 (*Dcr2* KO) and AF525 (*Ago2* KO). Cells were infected with ASALV (MOI 0.5). Total RNA was isolated at 48 hpi from the cells, small RNAs (18-40 nt) were sequenced and mapped to the ASALV genome (sense, positive numbers) and antigenome (antisense, negative numbers). **A**. Distribution of the small RNA lengths. Y-axis shows the proportion of small RNAs of a given length to total ASALV-specific small RNA reads. **B**. Mapping of 21nts and **C**. 27nts small RNAs across the ASALV genome and antigenome. The figure is a representative result of two independent experiments.

In AF5 cells, ASALV-specific siRNAs (Figure 3A) are produced and mapped across the genome (sense) and antigenome (antisense, Figure 3B), similar to the results previously observed in U4.4 cells (Hermanns et al., 2020). Similarly, in *Ago2* KO AF525 cells, the majority of ASALV-specific small RNAs are 21 nts long vsiRNAs (Figure 3A). They also map across the whole genome and antigenome, although with a bias to the 5’and 3’end (Figure 3B); which is not observed in AF5 cells. In *Dcr2* KO AF319, ASALV-specific siRNAs are strongly decreased, and a majority of them map to the 3’end of the ASALV genome.

piRNA-sized small RNAs were observed at a low concentration in both AF5 and AF525 cells (Figure 3A-C) and did not show the “ping-pong” amplification characteristics (Figure 4). In contrast, AF319 cells produce ASALV-specific piRNA-sized small RNAs (Figure 3A-C) with the ping-pong amplification characteristics (Figure 4). Antisense and sense piRNA-sized small RNAs showed a clear 10 nucleotides overlap. Adenine was the most frequent nucleotide on the 10th position of the sense piRNA-sized small RNA sequence, although the bias was not very strong. In antisense piRNA-sized small RNA sequences, uridine was the most frequent nucleotide at the first position (Figure 4).

**Figure 4:**
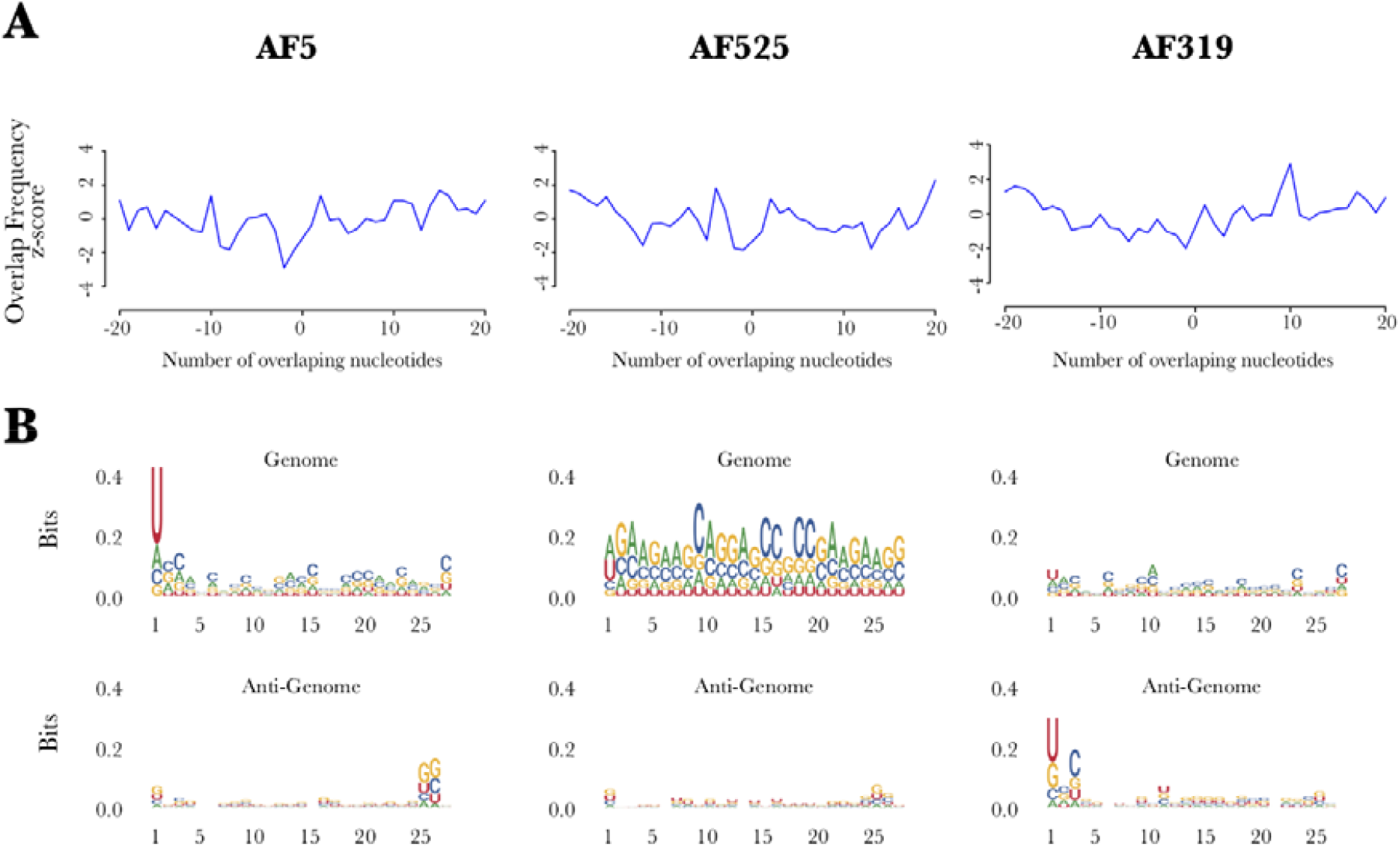
Characterization of ASALV specific 25-29 nts long small RNAs in *Ae. aegypti*-derived AF5, AF319 (*Dcr2* KO) and AF525 (*Ago2* KO). **A**. Overlap frequency of sense and antisense 25-29 nts long ASALV-specific small RNAs was calculated. **B**. Logo sequence plots show the sequence bias in various positions of 27 nts (as representative of vpiRNAs) long ASALV-specific small RNAs for genomic (upper panel) and antigenomic (lower panel) small RNAs. The figure is a representative result of two independent experiments.

In all cells, piRNA-sized small RNAs were mapped around the subgenomic promoter and 5’ end of the subgenomic RNA, encoding for the capsid protein, similar to vpiRNAs produced by arthropod-borne alphaviruses (Miesen et al., 2015; Schnettler et al., 2013). However, in AF319 cells, some piRNA-sized small RNAs map also to the 5’ of the genome (Figure 4C).

Taken together, vsiRNAs are the main small RNA species produced against ASALV infection under normal circumstances. In the absence of *Dcr2*, ASALV can induce piRNA-sized small RNAs with sequence characteristics indicative of the ping-pong amplification pathway.

### siRNA pathway, miRNA pathway and *Piwi4* are involved in the antiviral RNAi response against ASALV

Increased ASALV infection in the knock-out cell lines supports the involvement of the siRNA pathway in the antiviral defense against ASALV. To investigate the involvement of the other RNAi pathway proteins against ASALV in *Ae. aegypti-*derived AF5 cells, transcripts of different RNAi proteins were silenced by transfecting cells with sequence-specific dsRNAs (*Ago1, Ago2, Ago3, Piwi4, Piwi5, Piwi6*), prior to ASALV infection (MOI 0.5, Figure 5A).

**Figure 5:**
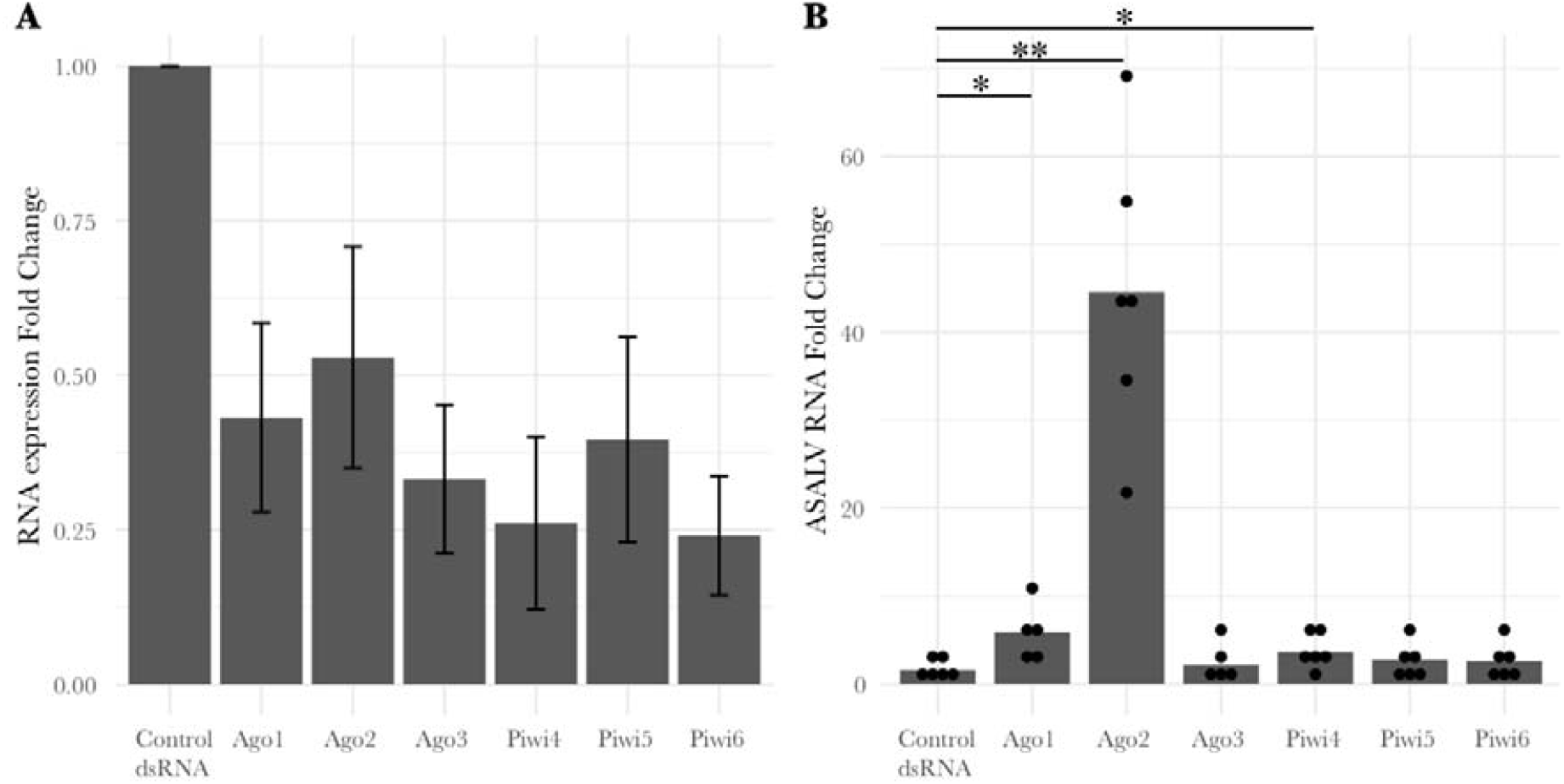
*Ago1, Ago2* and *Piwi4* silencing increases ASALV replication in *Ae*.. *aegypti* derived AF5 cells. Cells were transfected either with gene-specific dsRNAs or control dsRNA (LacZ-specific). The following day, cells were infected with ASALV (MOI 0.5), and total RNA was isolated 48 h post-infection. **A**. mRNA targets were quantified using gene-specific primers and Ribosomal protein S7 RNA as housekeeping transcript. 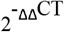 of mRNA targets was calculated with the mean normalised RNA expression of a given transcript in the control cells, within the same replicate, as control. The resulting mean fold change and standard error of the mean are shown. **B**. ASALV RNA was quantified using ASALV specific primers and Ribosomal protein S7 RNA as housekeeping transcript. ASALV RNA fold change was calculated using 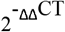 method with the mean of normalised expression of ASALV RNA, of all replicates, in the control cells as control. Bar plots represent the mean fold change for each group calculated. At least five independent replicates were performed. (*: p=<0.05, **: p<0.01).

Successful silencing was verified (Figure 5A), and viral RNA was quantified in the cells at 48 hpi and compared to control cells (transfected with dsRNA specific to LacZ). Viral replication increased significantly in cells where *Ago1* (t = 2.817, df = 4.665, p = 0.040), *Ago2* (t = 6.437, df = 5.039, p = 0.001) or *Piwi4* (t = 2.628, df = 8.543, p = 0.029) transcripts were silenced (Figure 5B). The ASALV RNA fold change was more pronounced in *Ago2* silenced cells than in *Ago1* and *Piwi4* silenced cells (Table S2). Furthermore, when *Piwi4* the silencing was conducted using Piwi4 siRNAs, instead of dsRNAs, ASALV replication increased although this increase was not significant (Figure S3).

### No RNAi suppressor effect of ASALV was detected in AF5 cells

Several insect viruses have been reported to encode proteins that interfere with the antiviral RNAi pathway, named viral suppressors of RNAi (VSR). VSRs can interfere at different steps of the RNAi pathways by interacting with key molecules (e.g. dsRNA or siRNAs) or proteins (e.g. *Ago2, Dcr2*), mostly of the exo-siRNA pathway. To determine if ASALV can suppress the exo-siRNA response in mosquito-derived cells, a previously used luciferase-based RNAi suppressor assay was performed (Ongus et al., 2006; Varjak, Maringer, et al., 2017). AF5 cells were either infected with ASALV (MOI 10) or mock-infected. After 24 hpi, cells were co-transfected with Firefly and *Renilla* luciferase (internal control) expression constructs as well as dsRNA (FFluc or LacZ as control) or siRNA (siFFluc, siHyg as control) to induce silencing. Luciferase activity was measured 24 hpi and sequence-specific silencing of Firefly luciferase in ASALV or mock-infected cells were compared.

Relative luciferase activity was significantly reduced in cells transfected with FFluc dsRNA compared to controls in both ASALV (t = -12.785, df = 2, p = 0.006; Figure 6A) and mock-infected cells (t = - 65.212, df = 2, p < 0.001; Figure 6A). Similarly, luciferase expression was significantly silenced when ASALV-infected (t = -15.469, df = 2, p= 0.004) or mock -infected cells (t = -20.322, df = 2, p= 0.002) were transfected with siFFluc compared to control siHyg transfection (Figure 6B). No difference in silencing of luciferase could be observed between mock or ASALV infected cells whether the silencing was induced by dsRNA (t = 0.281, df = 2.160, p = 0.803; Figure6A) or siRNA (t = 0.881, df = 4, p = 0.428; Figure 6B). Hence in our experimental setting, we did not detect any significant RNAi suppressor activity of ASALV in AF5 cells.

**Figure 6:**
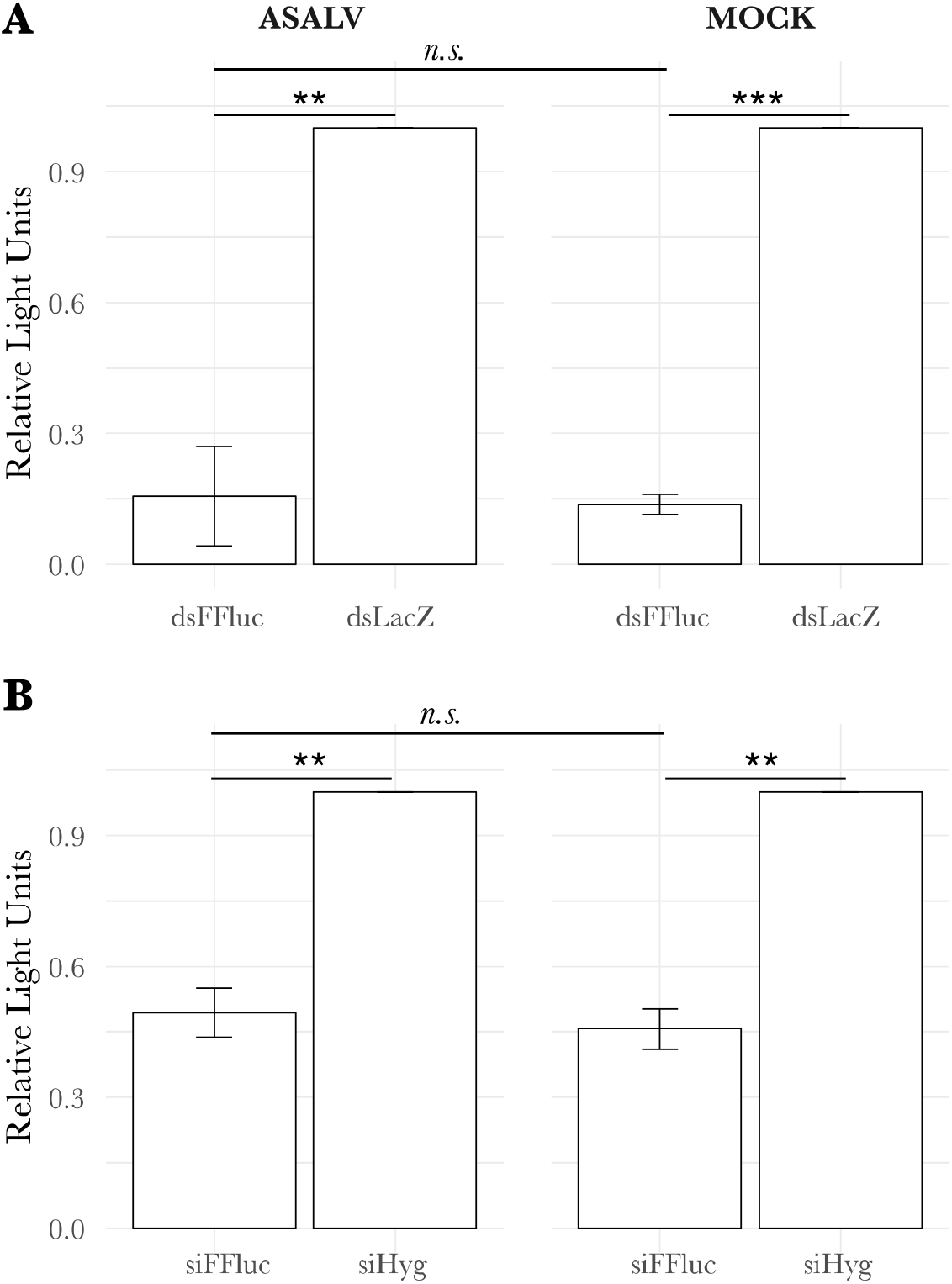
No RNAi suppressor effect of ASALV was detected in AF5 cells. AF5 cells are either mock-infected (cell culture media) or infected with ASALV (MOI 10). Following, cells were transfected with Firefly (FFluc) and *Renilla* luciferase (*Rluc*) expression constructs and either with 0.5 ng dsRNA (**A**) or 0.1 ng siRNA (**B**). Luciferase was measured using the Dual luciferase assay and FFluc expression was normalised to *Rluc* as an internal control (relative light units). FFluc/<u>Rluc</u> expression in the dsRNA(dsFluc) or siRNA(siFluc) transfected cells were normalised to control transfected cells (dsLacZ or siHyg). The mean of three independent experiments in triplicates are shown with SEM (***: p<0.001, **: p<0.01, n.s.: not significant).

## Discussion

RNA interference (RNAi) is an important antiviral response in insects, including mosquitoes. The interaction between the mosquito RNAi pathways and a variety of viruses can be identified by detecting virus-specific small RNAs and increased viral infection in case of silencing of key proteins of the different RNAi pathways. RNAi has been shown to act antiviral in mosquitoes against all tested viruses so far, although differences regarding the importance of specific pathways or proteins have been reported (Liu et al., 2019). Our knowledge about the antiviral RNAi response in mosquitoes comes from arbovirus studies, although mosquitoes often harbour insect-specific viruses (ISVs). Small RNAs specific to a variety of ISVs were found in infected cells and mosquitoes. However, the antiviral role of the RNAi pathway against ISVs is not known (Agboli et al., 2019). Here we identified the antiviral function of the mosquito RNAi pathways against an insect-specific alphavirus for the first time.

The only previous study investigating an RNAi response specific to an insect-specific alphavirus showed the production of ASALV-specific 21 nts vsiRNAs in *Ae. albopictus* derived (U4.4) cells; although no vpiRNAs was observed (Hermanns et al., 2020). Our results confirm this previously reported lack of ASALV-specific piRNA production in *Ae. aegypti*-derived RNAi competent AF5 cells (Figure 4). In contrast, arthropod-borne alphaviruses induce both vsiRNAs and vpiRNAs *in vitro* in *Ae. aegypti* and *Ae. albopictus*-derived cell lines, as well as in mosquitoes (Cirimotich et al., 2009; Goic et al., 2016; Morazzani et al., 2012; Schnettler et al., 2013; Siu et al., 2011a; Vodovar et al., 2012). Despite the difference in the small RNAs that are produced during infection, the mapping of ASALV specific siRNAs (both in AF5 and U4.4 cells) was very similar to the mapping of arthropod-borne alphaviruses. Both map along the genome and antigenome, more or less equally with some cold and hot spots (Morazzani et al., 2012; Schnettler et al., 2013; Siu et al., 2011b). This suggests that similar to arthropod-borne alphaviruses, ASALV also mainly induces vsiRNA production through dsRNA replicative intermediates.

ASALV replication is increased in both *Ago2* silenced (Figure 5B, Table S2) and *Ago2* or *Dcr2* knock-out cells (Figure 2); highlighting the antiviral role of the exo-siRNA pathway against ASALV. Similarly, silencing or knock-out of *Ago2* or *Dcr2* induced an increase in infection of tested arthropod-borne alphaviruses (Campbell et al., 2008; Schnettler et al., 2013; Sucupira et al., 2020; Varjak, Dietrich, et al., 2018; Varjak, Donald, et al., 2017). Furthermore, similar results have been found for arboviruses belonging to other virus families or orders (Liu et al., 2019), except for ZIKV, where no antiviral activity was reported for *Ago2* (Scherer et al., 2021; Varjak, Donald, et al., 2017). For the arthropod-borne alphavirus SFV, the magnitude of increase in infection was similar in *Dcr2* and *Ago2* knock-out cells (Scherer et al., 2021). In contrast, for ASALV, the differences between *Ago2* and *Dcr2* knock-out cells suggest an additional role of *Dcr2* in the antiviral response against ASALV independent of *Ago2*. For instance, *Dcr2* can detect viral RNA and induce an antiviral protein, Vago, which activates the Jak-STAT pathway leading to an antiviral effect in *Culex quinquefasciatus* (Hsu)-derived cells (Paradkar et al., 2012, 2014). Notably, however, Vago does not seem to be induced in infected *Ae. aegypti*-derived Aag2 cells (Russell et al., 2021). Alternatively, this increased antiviral effect of *Dcr2* against ASALV might be linked to another yet unknown antiviral pathway related to *Dcr2* activity.

ASALV-specific piRNA-sized small RNAs with ping-pong amplification characteristics were produced only in *Dcr2* knock-out cells. Previous reports have also shown an increase of SFV-specific vpiRNAs in cells lacking the *Dcr2* protein (Varjak, Maringer, et al., 2017b). It is possible that the increase in the vpiRNA production is a result of (i) the increased viral replication due to the lack of the antiviral *Dcr2* protein, (ii) the high concentration of ASALV RNA in the cytoplasm that is not cut into vsiRNAs or (iii) a combination of both. Although ASALV replication was increased in *Ago2* knock-out cells, no ping-pong specific vpiRNAs were detected. While this could mean that increased viral replication is not solely sufficient for ASALV specific vpiRNA production, it has to be noted that the increase in ASALV replication in *Ago2* knock-out cells was still lower compared to *Dcr2* knock-out cells. Therefore, it could be that the increased ASALV RNA concentration in *Ago2* KO cells is not sufficient to trigger vpiRNA production, in contrast to *Dcr2* KO cells. In addition, it is likely that in *Dcr2* knock-out cells specifically, the amount of viral dsRNA molecules would increase. As the precise trigger for vpiRNA production in mosquitoes is not yet known, it could be that the concentration of ASALV dsRNA in *Dcr2* knock-out cells could play a role in triggering vpiRNA production. On the other hand, the putative essential proteins for the biogenesis of vpiRNAs, *Piwi5* and *Ago3*, were not antiviral against ASALV (Figure 5B), consistent with findings from arthropod-borne alphaviruses(Miesen et al., 2015; Varjak, Dietrich, et al., 2018; Varjak, Donald, et al., 2017).

Silencing of *Piwi4* resulted in a small but significant increase in ASALV replication as it has previously been shown for other arboviruses, including alphaviruses (Dietrich, Shi, et al., 2017; Varjak, Maringer, et al., 2017b). The general antiviral role of *Piwi4* is still not clear. Piwi4 is not required for the production of SFV- or SINV-specific vpiRNAs, but it has recently been shown to bind DENV-specific piRNAs derived from viral cDNA in infected *Ae. aegypti* (Tassetto et al., 2019). While an interaction between *Piwi4* and piRNA as well as siRNA pathway proteins, including *Dcr2*, has previously been shown, *Piwi4* antiviral activity is independent of *Dcr2* in SFV infected cells (Joosten et al., 2021; Varjak, Maringer, et al., 2017a). To check this for ASALV, we silenced *Piwi4* by adding siRNAs, both in *Dcr2* competent and knock-out cell lines. While the silencing of *Piwi4* through siRNA increased ASALV replication, the increase was not significant in either of the cell lines (Figure S3). If this is due to the slightly lower silencing efficiency with siRNAs compared to dsRNA is not known. Hence it was not possible to conclude whether the effect of *Piwi4* is *Dcr2* independent.

Our results suggest an antiviral effect of *Ago1*, which is primarily involved in the miRNA pathway (Figure 5). Although the mosquito miRNA response has been shown to interact with viruses through either mosquito or virus-encoded miRNAs (Leggewie & Schnettler, 2018), silencing of *Ago1* has not resulted in changes of arboviral alphavirus replication (Keene et al., 2004; McFarlane et al., 2014; Schnettler et al., 2013). Similar increases in virus infection upon Ago1 silencing have been reported for midge-borne orthobunyaviruses in *Ae. aegypti* derived cells in contrast to mosquito-borne orthobunyaviruses (Dietrich, Shi, et al., 2017). Additional experiments are needed to determine if the difference in *Ago1* activity against arthropod-borne alphaviruses compared to insect-specific alphaviruses can be generalised.

Many viruses infecting insects encode proteins to suppress the RNAi pathway, such as Flock House Virus or Culex Y virus (O’Neal et al., 2014). Several arboviruses, such as Dengue and West Nile Virus, have also been shown to interfere with the RNAi response by employing competitive substrates for Dcr2 derived from their nucleic acids (O’Neal et al., 2014). Furthermore, recent work has identified the non-structural protein NS2A of flaviviruses as a potent suppressor of RNAi (Qiu et al., 2020). In our experimental system, we did not observe any RNAi suppressor activity of ASALV.

ISVs belonging to some of the arbovirus families and orders, such as *Bunyavirales* (Marklewitz et al., 2015) and *Flaviviridae* (Cook et al., 2019), are thought to be ancestral to arboviruses, suggesting that dual-host (invertebrate-vertebrate) tropism evolved from invertebrate specific viruses. As not many insect-specific alphaviruses have been discovered so far, it is difficult to identify whether insect-specific viruses or the arthropod-borne alphaviruses are the ancestors in the alphavirus genus (Halbach et al., 2017). Nevertheless, like other insect-specific alphaviruses so far, ASALV is basal to the Western Equine encephalitis virus complex clade, suggesting arthropod-borne alphaviruses in this clade could have evolved from an ancestral insect-specific virus (Halbach et al., 2017; Hermanns et al., 2020). It is also possible that the changes in the mosquito-virus interactions drive their evolution resulting in their ability to transmit to vertebrates. In this context, differences between arboviral and insect-specific alphaviruses’ interaction with mosquito RNAi pathways could be one of the reasons why ISVs were restricted to invertebrate hosts. In contrast to arthropod-borne alphaviruses studied so far, we showed that ASALV specific vpiRNAs are not produced in *Dcr2* competent cells, and *Ago1* was antiviral against ASALV. However, to be able to generalise this observation to other insect-specific alphaviruses, more studies describing their interactions with mosquito hosts are needed. Further studies taking both the persistent nature of ISVs and the tissue-specificity of the RNAi response into account could determine whether the interactions of insect-specific alphaviruses with the RNAi pathways restrict ISVs to their mosquito hosts.

