## Supplementary Material for "Antiviral RNAi response against the insect-specific Agua Salud alphavirus"

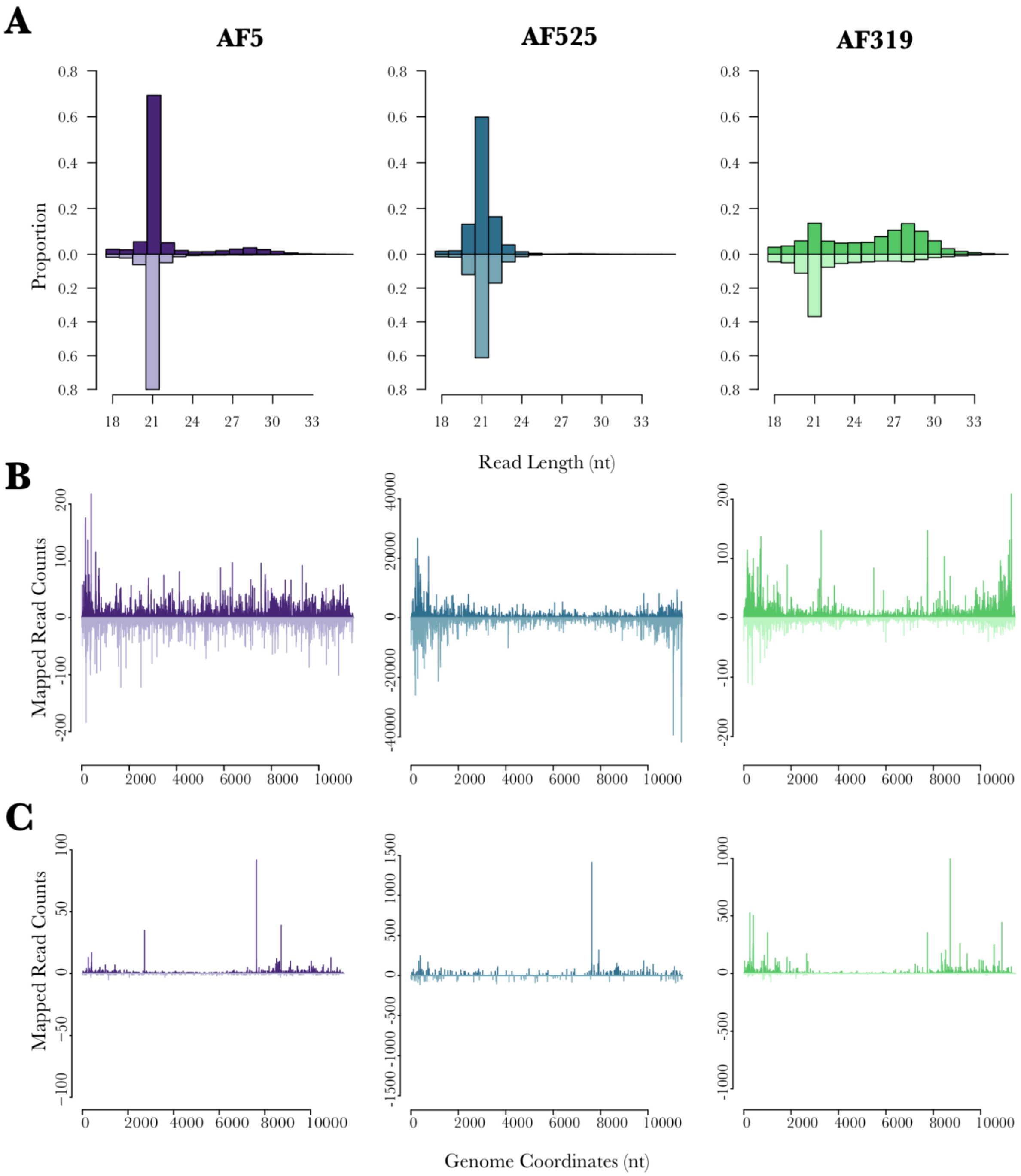


**Figure S1: ASALV-specific small RNA production in *Ae. aegypti*-derived AF5, AF319 (Dcr2 KO) and AF525 (Ago2 KO).** Cells were infected with ASALV (MOI 0.5). Total RNA was isolated at 48 hpi from the cells, small RNAs (18-40 nt) were sequenced and mapped to ASALV genome (sense, positive numbers) and antigenome (antisense, negative numbers). A. Distribution of the small RNA length. Y axis shows the proportion of small RNAs of a given length to total ASALV-specific small RNA reads. B. Mapping of 21 nts and C. 27 nts small RNAs on ASALV genome and antigenome. Representative of two repeats.

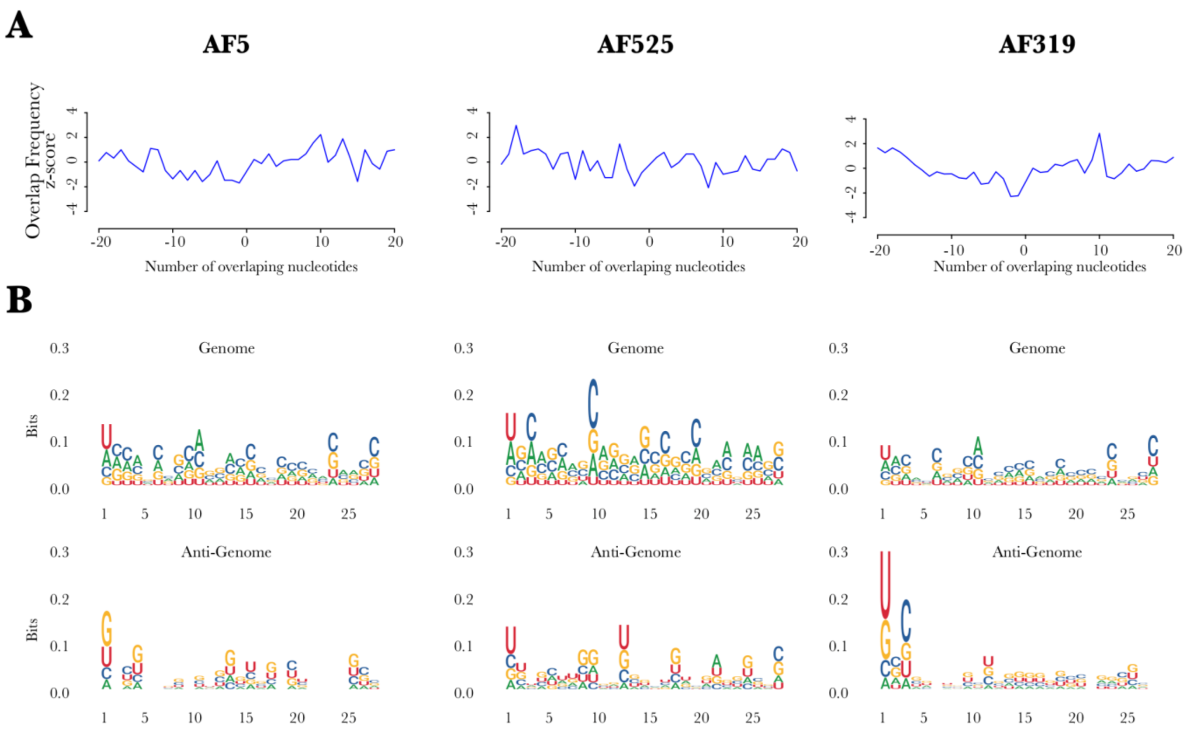


**Figure S2 Characterization of ASALV specific 25-29 nts long small RNAs in *Ae. aegypti*-derived AF5, AF319 (Dcr2 KO) and AF525 (Ago2 KO). A.** Overlap frequency of sense and anti-sense 25-29 nts long small RNAs is calculated. **B.** Logo sequence plots show the sequence bias in various positions of 27 nts (as representative of vpiRNAs) long ASALV-specific small RNAs, genome (upper panel), antigenome (lower panel). Representative of two repeats.

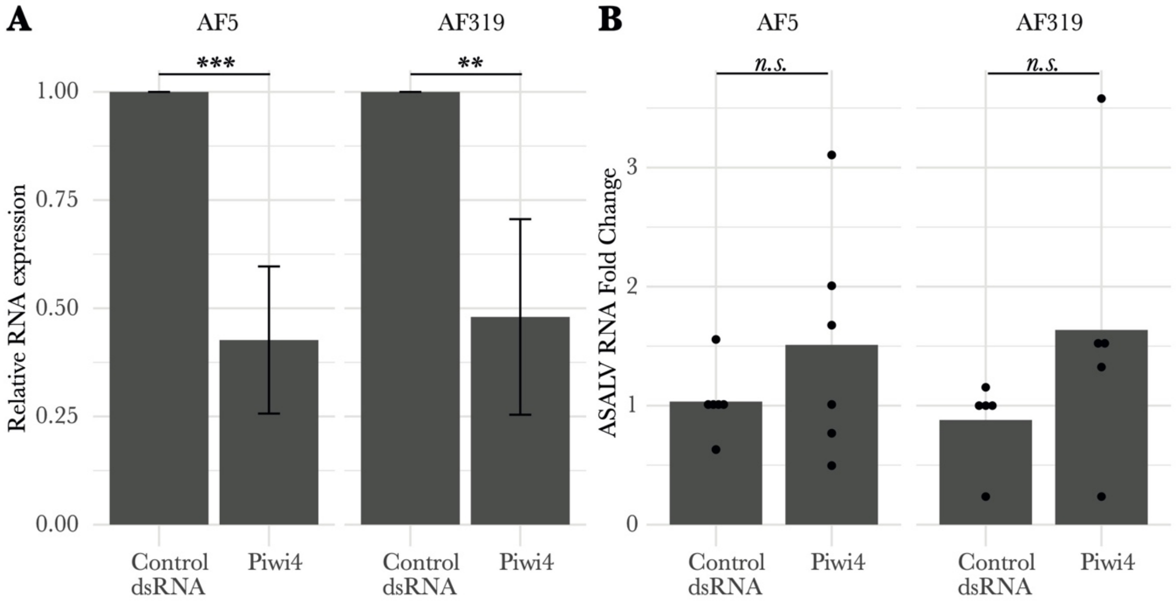


**Figure S3: *Piwi4* knock-down in Ae. aegypti-derived *Dcr2* KO AF319 cells. AF319** and AF5 cells were transfected with either Piwi4 specific siRNA or control siRNA, followed by ASALV (MOI 0.5) infection. *Piwi4* transcript (A) and ASALV RNA fold change (B) in infected cells were quantified at 48 hpi, using the $2^{-\Delta\Delta CT}$ method with Ribosomal protein S7 RNA as housekeeping gene and control siRNA transfected cells. Six independent replicates were performed for AF5 and AF319 cells. Bar plots represent the mean of the performed replicates (*: p=<0.05, **: p<0.01, ns: not significant).

**Table S1. qPCR primers used in the study.**

| | | |
|---|---|---|
| **Target** | **Forward/ Reverse primer (5' -3')** | **Reference** |
| Ago1 (*Aedes aegypti*) | GTACGATGCGTCGTAAGTAC/GTACTTGTCGAGGAAGTATTTGG | This study |

| | | |
|---|---|---|
| Ago2 (*Aedes aegypti*) | GGCTGCTCACCCAATGTATCAAGA/AACCGTTCGTTTTGGCGTTGAT | (1) |
| Piwi4 (*Aedes aegypti*) | CTTCTCCACCACAGCCAATG/GTCCAATCTGCCTGTTCTCCA | (1) |
| Ago3 (*Aedes aegypti*) | TGCTCCAGACGACGGTTTTG/GGGTCAATATAACGGCTCCCAG | This study |
| Piwi5 (*Aedes aegypti*) | CAGTTTTGGAAGACAGAGTTGGA/CCTGCCGTCACTTTGTAATTTTC | This study |
| Piwi6 (*Aedes aegypti*) | TCCGACGTTTTCAAGTTTTGGA / CACTTTACACTGATCCTGCTCG | This study |
| S7 (*Aedes aegypti*) | CCAGGCTATCCTGGAGTTG/ GACGTGCTTGCCGGAGAAC | (1) |

| | | |
|---|---|---|
| ASALV | CCGTACTCGAAACAGACATTGC/ TCGTCAACGCCTAGATCCTCTA | (2) |

1. M. Varjak, *et al.*, Characterization of the Zika virus induced small RNA response in Aedes aegypti cells. *PLoS Negl. Trop. Dis.* **11**, 1–18 (2017).

2. K. Hermanns, *et al.*, Agua Salud alphavirus defines a novel lineage of insect-specific alphaviruses discovered in the New World. *J. Gen. Virol.* **101**, 96–104 (2020).

**Table S2: ASALV RNA fold change in knockdown experiments.**

| mRNA Target | ASALV fold change | Standard deviation |
|---|---|---|
| Control | 1.609 | 1.024 |
| Ago1 | 5.868 | 3.248 |
| Ago2 | 44.569 | 16.315 |
| Ago3 | 2.143 | 2.061 |
| Piwi4 | 3.638 | 1.589 |
| Piwi5 | 2.762 | 2.321 |
| Piwi6 | 2.562 | 1.582 |

Values represent the mean of at least 5 replicates that were performed. Values are rounded to 3 decimal points.
